# Adaptive periodicity in the infectivity of malaria gametocytes to mosquitoes

**DOI:** 10.1101/294942

**Authors:** Petra Schneider, Samuel S. C. Rund, Natasha L. Smith, Kimberley F. Prior, Aidan J. O’Donnell, Sarah E. Reece

## Abstract

That periodicity in the biting activity of mosquito vectors explains why malaria parasites have evolved rhythms in cycles of asexual replication in the host’s blood was proposed almost 50 years ago. Yet, tests of this hypothesis have proved inconclusive. Using the rodent malaria *Plasmodium chabaudi*, we examine rhythms in the density and infectivity of transmission forms (gametocytes) in the host’s blood, parasite development inside mosquitoes, and onwards transmission.Moreover, we control for the confounding effects of rhythms in mosquito susceptibility. We reveal that at night, gametocytes are twice as infective to mosquitoes, despite being less numerous in the blood. This enhanced infectiousness at night interacts with mosquito rhythms to increase sporozoite burdens by almost four-fold when mosquitoes feed during their day. Thus, daytime blood-feeding (*e.g.* driven by the use of bed nets) may render gametocytes less infective, but this is compensated for by the greater susceptibility of mosquitoes.

## Introduction

The rotation of the Earth, every 24-hours, results in exposure to environmental (abiotic) rhythms in ambient light, temperature, humidity, and UV radiation. The evolution of circadian rhythms is assumed to be an adaptation to cope with the challenges of - and exploit the opportunities provided by - predictable environmental rhythms [1,2]. Parasites also experience diverse environmental rhythms generated by daily rhythms in their hosts (and vectors) [3–5]. Such “biotic” environmental rhythms include immune responses, resource availability, and transmission opportunities. For example, Hawking observed that the microfilaria (transmission forms) of *Wuchereria bancrofti* transmitted by nocturnally active mosquitoes are only present in the peripheral blood of the host during the evening, migrating to the lungs during the day [6]. The timing of this migration behaviour is reversed for the Pacific type of *W. bancrofti* which is transmitted by a diurnal vector species [6,7].

The ability to schedule the expression of transmission traits for the time-of-day that vectors are host-seeking appears to be adaptive (*i.e.* maximises parasite fitness) and could benefit many species of parasites. Thus, Hawking also predicted this hypothesis explains periodicity in the cycles of asexual replication observed in many species of malaria parasite [8]. In the blood of the vertebrate host, malaria (*Plasmodium*) species undergo successive, synchronised, cycles of asexual replication which result in bouts of fever every 24, 48, or 72 hours (depending on the species) when parasites burst to release their progeny. The regularity of fevers is sufficiently reliable that it was used as a diagnostic symptom in the Hippocratic era, but why there appears to be a circadian basis to the duration of asexual cycles has been a mystery ever since [3,9,10]. Hawking proposed that malaria parasites burst to release their progeny (schizogony) at a specific time of day to coincide the maturation of transmissible forms (gametocytes) with the nocturnal foraging activity of their mosquito vectors [6,8].

Tests of whether Hawking’s hypothesis applies to malaria parasites have so far proved inconclusive [5]. Hawking himself observed that mosquitoes feeding at night harboured more *P. knowlesi* oocysts [8]. But Karunaweera noted that mosquitoes fed at night time, during the height of *P. vivax* fever, produce less oocysts then mosquitoes fed earlier in the day [11]; no influence of time-of-day on oocyst burden has been reported for *P. chabaudi* [12] nor *P. falciparum* [13,14]; and gametocyte density can peak in the blood at the opposite time-of-day to when mosquitoes forage [15]. However, some of these studies were not well replicated, with fewer than three mosquitoes observed per time point. Despite the seeming lack of support, it is important not to prematurely reject Hawking’s hypothesis. Previous studies have neglected to control for circadian rhythms in mosquito physiologies, including immune responses [16,17]. If rhythms exist in parasite transmissibility and mosquito susceptibility to infection, but their timing is inverted, then times-of-day when parasites are most transmissible are opposed by reduced vector susceptibility. In this case, the net effect appears to erode rhythms in both parasite transmissibility and mosquito susceptibility. In this case, the net effect will dampen rhythms in both parasite transmissibility and mosquito resistance. Furthermore, studies that only compare two time points of a single rhythm may suffer from coincidently sampling when the rhythm reaches the same trait value as it ascends and descends (particularly if studying time points are 12 hours apart on a sine wave).

Understanding how rhythms in gametocyte infectivity interact with mosquito rhythms is urgently needed because mosquito populations are responding to the use of bednets by shifting the time-of-day they forage for blood [18]. Here, we find support for Hawking’s hypothesis by independently testing the roles of both parasite rhythms and vector rhythms, and examining parasite transmissibility from the host’s blood throughout development in the vector. Specifically, we manipulate the time-of-day of blood feeding for both the mosquito vector *Anopheles stephensi* and the rodent malaria parasite *P. chabaudi*, and we quantify the density of mature gametocytes in the blood at the time of biting, as well as intensities of the resulting oocyst and sporozoite infections in the vector. We find that at night time, despite being less numerous in the blood, gametocytes are twice as infective to mosquitoes. The enhanced infectiousness of gametocytes at night interacts with greater mosquito susceptibility in the daytime to elevate sporozoite burdens by almost four-fold.

## Results

Our experiment involved four treatment groups in which mosquitoes experiencing their daytime or night time fed on malaria-infected mice experiencing their own daytime or night time (Figure 1). For both parasites and mosquitoes, daytime refers to Zeitgeber Time 8 (ZT; hours since lights on) and night time refers to ZT16. We chose these times because ZT8 falls in the middle of the mosquito’s resting phase and at ZT16, they are several hours into their period of nocturnal activity (SI Figure 1). Each treatment group comprised of 20 mice, resulting in 80 mosquito feeds (80 cages of 60 mosquitoes/cage) across the whole experiment and totalling 2400 mosquitoes. Blood stage *P. chabaudi* parasites have a 24-hour rhythm in their cycles of asexual replication and are early in the developmental cycle (young trophozoites) at ZT8 and approaching maturation (schizogony) at ZT16 [19]. At the time of each feed, we quantified gametocytes by RT-qPCR (because this method detects mature gametocytes), before exposing each mouse to a cage of female mosquitoes. We subsequently quantified the parasite burdens on the midguts (oocysts) and in the salivary glands (sporozoites) as well the prevalence of infections.

**Figure 1.**
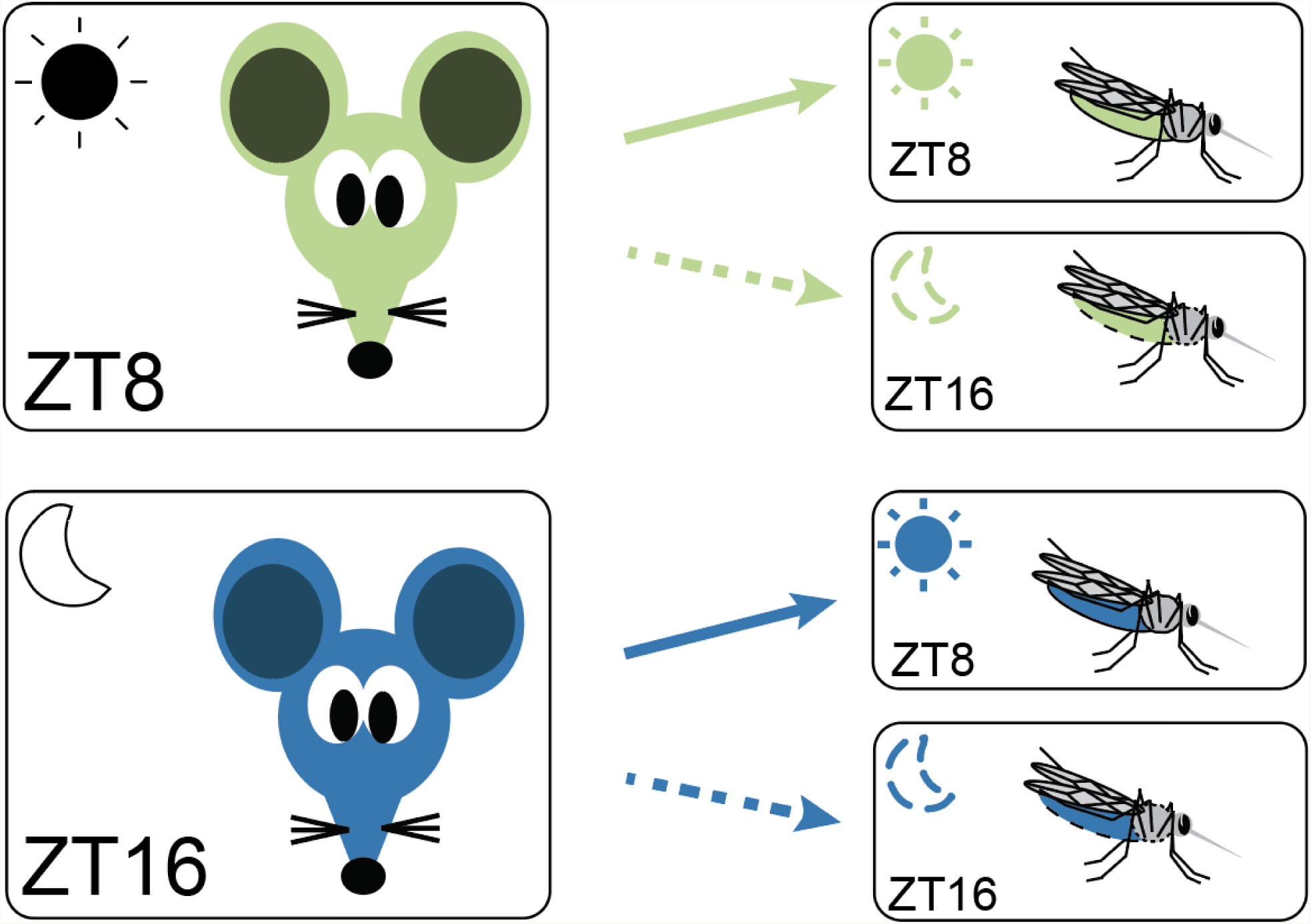
Experimental design. We infected 80 mice with ring stages of *P. chabaudi* (genotype AS) at ZT3 to ensure parasite rhythms were in phase with the host’s rhythms in all treatment groups [19,20]. At ZT8 (green) 40 mice were fed to 40 cages of mosquitoes experiencing their day (ZT8; solid arrow) or night (ZT16; dotted arrow) and we repeated this at ZT16 for the other 40 mice (blue). Gametocyte metrics were quantified in the two groups of mice just before exposure to mosquitoes and oocysts and sporozoites were followed in the four groups of mosquitoes. All feeds (*i.e.* at both ZT8 and ZT16) were performed in the dark to prevent unexpected light exposure to mosquitoes, which is known to affect biting behaviours and rhythms in gene expression [21,22].

### Fewer gametocytes circulate at night

We first examined whether the densities of gametocytes circulating in host blood differs between day and night. Phenomena such as timed release from the bone marrow and age-specific mortality rates of gametocytes have been proposed to generate daily rhythms in the availability of gametocytes to mosquitoes [23]. Hawking’s hypothesis would be supported if parasites schedule their development to maximise the appearance of gametocytes in the blood at night. However, we find that the densities of mature gametocytes are on average 1.6-fold (± SEM 0.2) lower at night compared to the day (F_1,78_= 11.11, *P*=0.001) (Figure 2). We confirmed this result in an independent experiment using a different *P. chabaudi* genotype (genotype CR), finding that gametocyte density is 1.5-fold (± SEM 0.2) lower on average at night (ZT17-21) compared to the day (ZT5-9) (F_1,14_=5.70, *P*=0.032).

**Figure 2.**
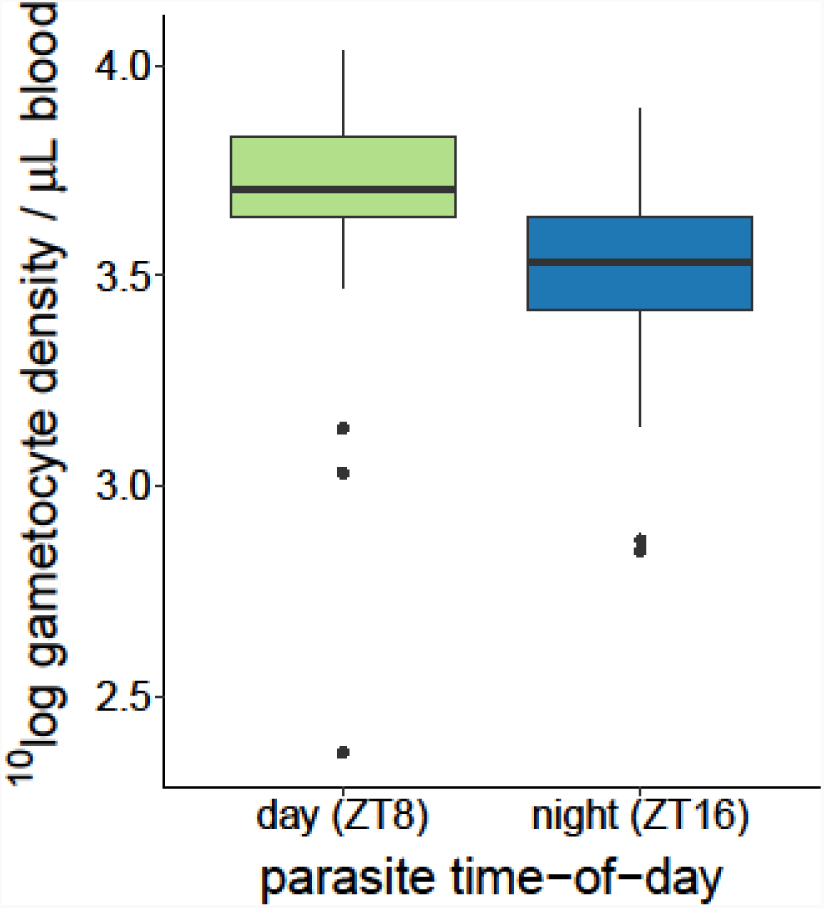
Gametocyte densities circulating in host blood are lower during the night time (ZT16) than the daytime (ZT8). N=20 mice per group.

Blood samples to quantify gametocytes are taken from the host’s tail vein, but mosquitoes harvest blood from subdermal capillaries. This raises the possibility that the lower density of gametocytes observed at night is a consequence of time-of-day specific accumulation in the capillaries [12,24,25]. Therefore, we carried out another experiment to simultaneously compare gametocyte densities in blood from the host’s tail vein to those in mosquito blood meals, at night (ZT16). When normalised to the density of white blood cells (following [25]), gametocyte densities varied across mice but are not significantly different in venous blood (0.62 ± SEM 0.19 gametocytes/white blood cell) and mosquito blood meals (0.57 ± SEM 0.15 gametocytes/white blood cell) (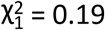, *P*=0.659). Thus, the lower density of gametocytes in host blood at ZT16 is not explained by their migration to the peripheral circulation.

### Mosquitoes are more likely to be infected from daytime blood meals

We observed no significant differences in the proportion of mosquitoes feeding (>93% fed in all cages) with respect to the time-of-day of feeding for parasites (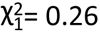, *P*=0.608), mosquitoes (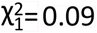, *P*=0.766), or their interaction (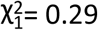, *P*=0.590). Night fed mosquitoes are 1.18-fold (± SEM 0.05) less likely to harbour oocysts compared to mosquitoes that fed in their daytime (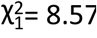, *P*=0.003), irrespective of parasite time (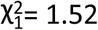, *P*=0.218), or the interaction between mosquito and parasite time (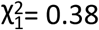, *P*=0.537) (Figure 3A). However, the oocyst burdens of infected mosquitoes are not significantly influenced by time-of-day of feeding for parasites (F_1,73_=2.46, *P*=0.121), mosquitoes (F_1,72_=1.19, *P*=0.280) or their interaction (F_1,68_=0.01, *P*=0.928) (Figure 3B).

**Figure 3.**
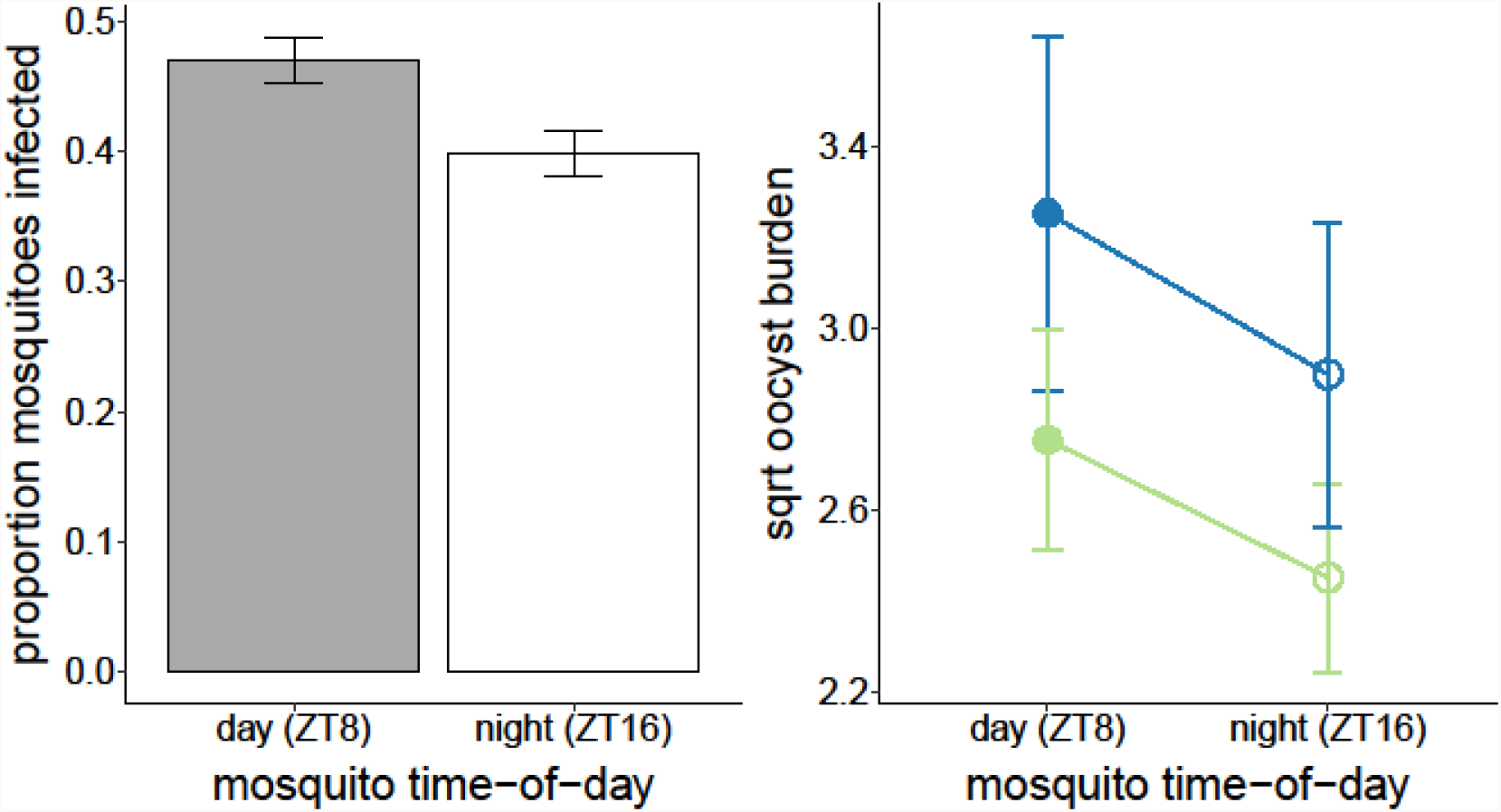
Night fed mosquitoes are less likely to be infected (A) but oocyst burdens are not influenced by time-of-day (B). Mean ± SEM proportion of mosquitoes that are infected with oocysts for daytime (ZT8; filled bar) and night time (ZT16; open bar) fed mosquitoes (A). Mean ± SEM oocyst burdens for mosquitoes that became infected after feeding on mice experiencing their day (ZT8; green) or night (ZT16; blue) by mosquitoes experiencing their day (ZT8; closed symbols) or night (ZT16; open symbols); square root transformed to meet model assumptions (B). Model estimates are plotted and a qualitatively similar pattern is observed for B when uninfected mosquitoes are included in the analysis.

### Gametocytes are more infectious at night

Fewer gametocytes are available to mosquitoes at night (Figure 2), yet this reduced density does not affect the prevalence or intensity of oocysts. This suggests gametocytes are more infectious at night. Infectivity can be assessed from the positive correlation between gametocyte density and oocyst burden; the steeper the slope the more likely a gametocyte is to develop into an oocyst. We find that time-of-day for mosquitoes (F_1,71_=1.20, *P*=0.277) and its interaction with parasite time-of-day (F_1,69_=0.02, *P*=0.887) do not significantly influence gametocyte infectiveness, but parasite time-of-day does (F_1,72_=18.92, *P*<0.001) (Figure 4). Specifically, each gametocyte, on average, results in 2.13-fold (± SEM 0.31) more oocysts when taken up at night compared to the daytime.

**Figure 4.**
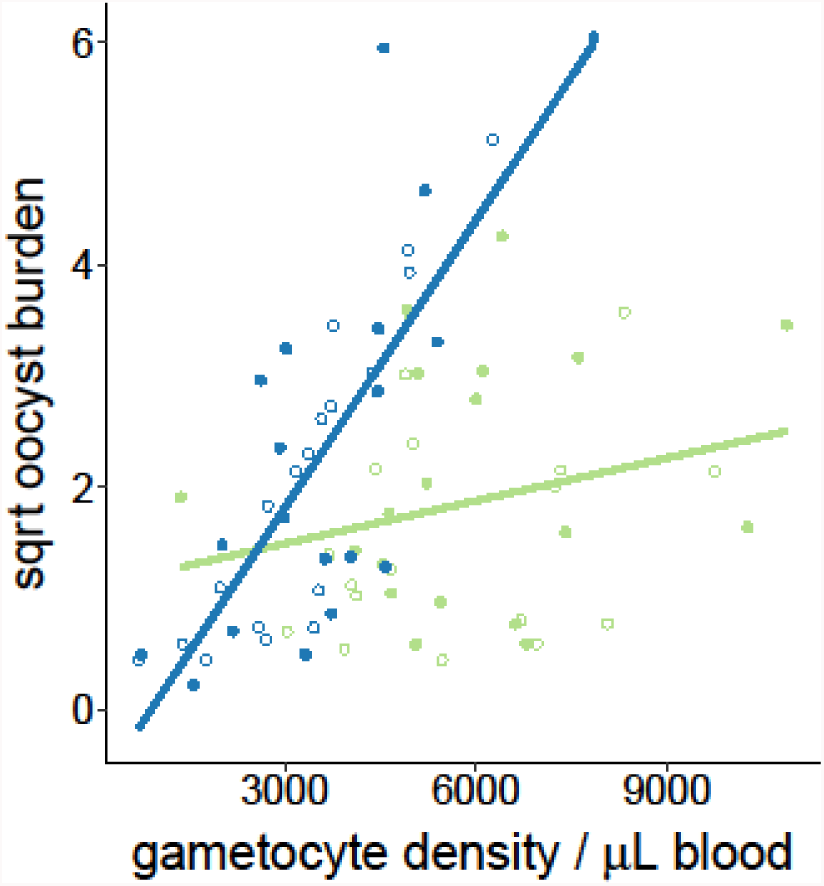
Gametocytes are more infective at night. Gametocytes taken up from hosts experiencing their night (ZT16; blue) are more likely to form oocysts than those taken up during the daytime (ZT8; green), regardless of time-of-day for mosquitoes (ZT8 closed, ZT16 open symbols). Gametocyte densities for each host are plotted against their corresponding mean oocyst burdens (square root transformed to meet model assumptions), and the fits are from linear regressions.

### Onwards transmission is determined by time-of-day of both parasites and mosquitoes

Hawking focussed on whether parasites have evolved to coordinate the maturation of gametocytes in the host’s blood with the blood-foraging activity of mosquito vectors. Yet, for such a strategy to be adaptive (*i.e.* maximise fitness), it should enhance the potential for transmission from vectors to new hosts. Only sporozoites in the salivary glands can be transmitted to new hosts so both the probability of a mosquito harbouring sporozoites and sporozoite density determine the likelihood of onwards transmission. Sporozoite prevalence did not vary with mosquito time-of-day (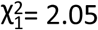, *P*=0.152; Figure 5A), parasite time-of-day (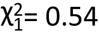, *P*=0.464) or their interaction (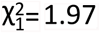, *P*=0.161), and thus does not reflect the lower prevalence in night-fed mosquitoes that we observed for oocysts (Figure 3A). However, sporozoite burdens are shaped by an interaction between time-of-day for both gametocytes and mosquitoes (F_1,73_=7.11, *P*=0.009 Figure 5B).Specifically, in mosquito samples harbouring sporozoites, burdens are 3.91-fold (± SEM 1.58) higher in mosquitoes that fed during their daytime, but only when taking up gametocytes experiencing their night time.

**Figure 5.**
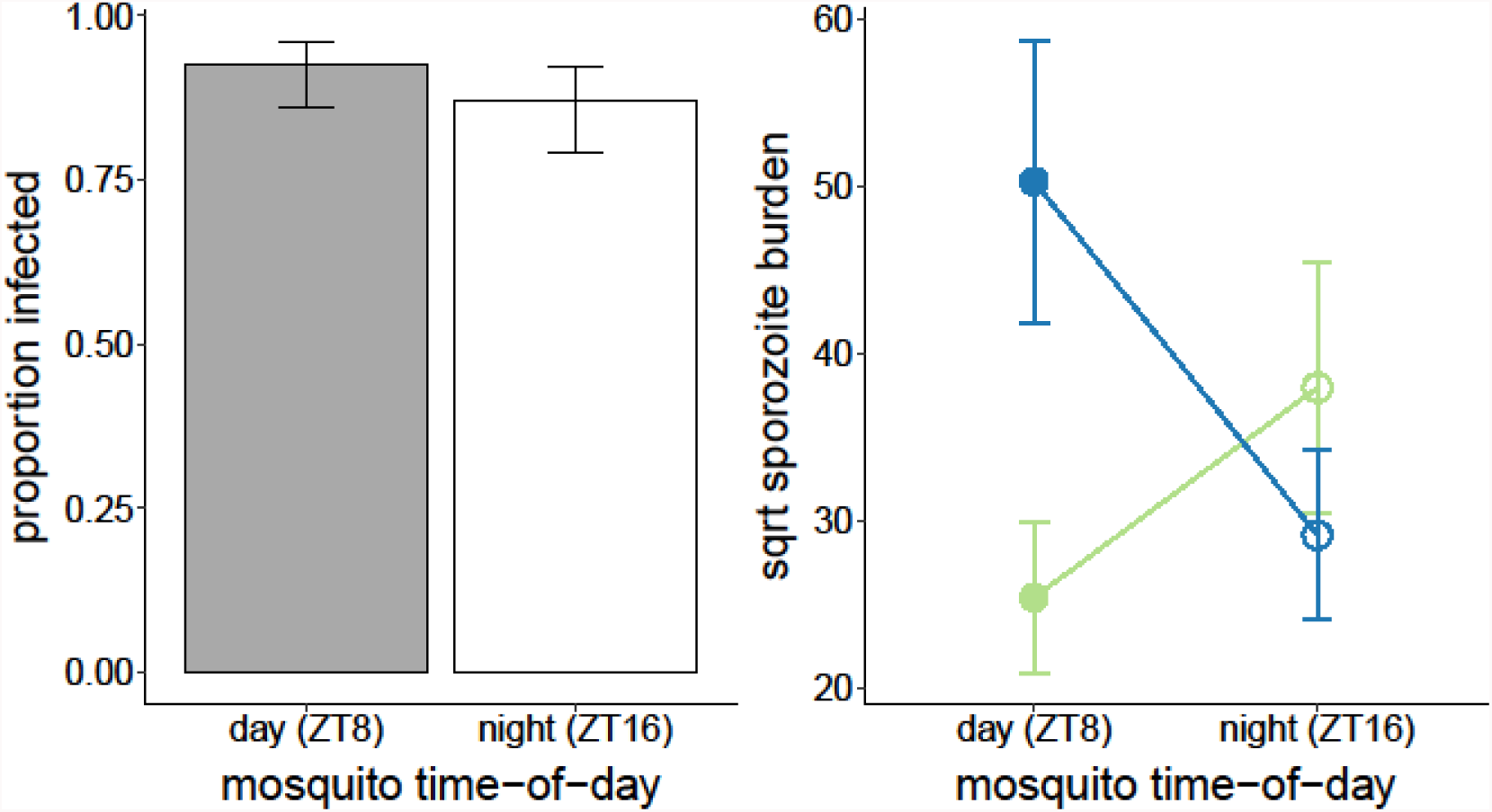
Parasite and mosquito time-of-day do not affect sporozoite prevalence (A) but do affect sporozoite burdens (B) Each sample consisted of a pool of 5 mosquitoes that blood fed on the same mouse (4 samples per mouse): a positive sample requires that at least 1 of 5 mosquitoes were infected with sporozoites. The mean ± SEM proportion of sporozoite positive samples for day and night feeding mosquitoes (A). Mean ± SEM sporozoite burdens for samples containing infected mosquitoes after feeding on mice experiencing their day (ZT8; green) or night (ZT16; blue) by mosquitoes experiencing their day (ZT8; closed symbols) or night (ZT16; open symbols); square root transformed to meet model assumptions (B). A qualitatively similar pattern is observed for B when samples containing only uninfected mosquitoes are included in the analysis as well.

## Discussion

We have deployed modern techniques to quantify gametocytes and assess malaria transmission in a well-replicated experiment to determine whether parasites maximise transmission by coordinating their development with the foraging activity of the mosquito vector. By separately assessing the effects of time-of-day on parasites and vectors we reveal that: (1) gametocytes are less numerous in the host’s blood at night but night time gametocytes are more likely to develop to oocysts; (2) the greater infectivity of night time gametocytes does not increase the probability that mosquitoes become infected or result in more intense infection at the oocyst stage; (3) independently of parasite time-of-day, mosquitoes fed at night are less likely to be infected with oocysts; and (4) the greater infectivity of night time gametocytes does not alter the probability of mosquitoes harbouring sporozoites but does increase sporozoite burden in mosquitoes fed in their daytime. We have previously shown that the timing of schizogony brings fitness benefits in the form of enhancing within-host survival [19], and now reveal that selection driven by periodicity in transmission to vectors can shape the timing of schizogony. However, any fitness benefits accrued by orienting the timing of schizogony are dependent on mosquito rhythms.

Finding that gametocytes are more infective at night supports Hawking’s hypothesis. Why might gametocytes be less numerous but more infective at night? Having ruled out accumulation in the peripheral capillaries at night, several non-mutually exclusive explanations remain. The developmental cycle of gametocytes offers a proximate (mechanistic) explanation (Figure 6). Gametocytes have a finite lifespan and host immune responses can clear circulating gametocytes [23]. Gametocytes quantified at ZT8 are the cohort (B, purple arrow) just reaching sexual maturity plus the survivors of the previously produced cohort (A, grey arrow), whereas gametocytes quantified at ZT16 comprise mature gametocytes from cohort B (purple arrow) plus even fewer survivors from cohort A (grey arrow). Thus, the loss of gametocytes as the circadian cycle progresses is an unavoidable consequence of extrinsic mortality, but the attainment of maturity of the next cohort compensates for the loss in numbers. In other words, if there was no enhancement in infectivity at night, parasites would be fitter if they transmitted during the daytime when gametocytes are more numerous.

**Figure 6.**
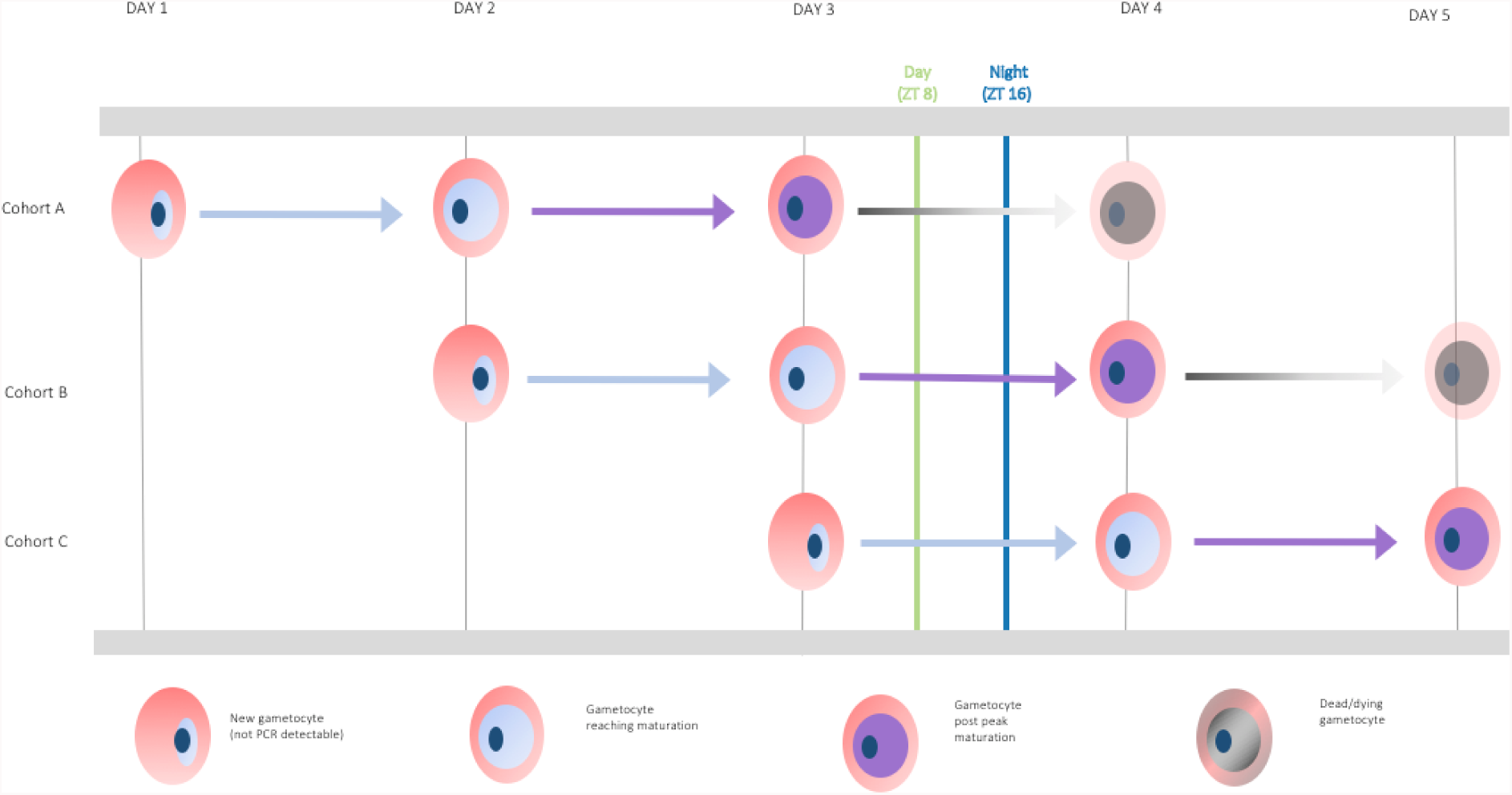
Dynamics of gametocyte development. A new cohort of gametocytes starts developing at every schizogony (~ZT17) and *P. chabaudi* gametocytes are thought to require approx. 36 hours to reach maturity, remain mature (*i.e.* infective) for 12-18 hours before senescing, and most gametocytes are cleared by 60 hours [12,26]. During our experiment we collected parasites at daytime (ZT8; green line) and night time (ZT16; blue line), thus sampling 3 cohorts of gametocytes (A-C) and our RT-qPCR assays detect gametocytes once they are mature (cohorts A and B). At ZT16 the bulk of the gametocytes present are infective because more of the senesced cohort from day 1 will have been cleared (indicated by the shading of the grey arrow) compared to at ZT8. At ZT8, sampled gametocytes comprised those produced on day 3 and not yet infective and not detected (cohort C); those produced on day 2 and at peak maturity (cohort B); and produced on day 1 and senescing (cohort A). Whereas, At ZT16, gametocytes were not yet infective and not detected (cohort C); at peak maturity (cohort B); and mostly cleared (cohort A).

Evolutionary (ultimate) explanations for timing gametocyte production (*i.e.* schizogony) according to Figure 6 may include maximising the proportion of gametocytes that are mature at the time of transmission. This metric will be under stronger selection than the density of mature gametocytes if immature or senesced gametocytes interfere with mating success in the blood meal. For instance, the time available to mate in the blood meal is limited and males are often limiting [27,28], and so, attempts to fertilise senesced females may “waste” males [29]. In this scenario, parasite evolution is shaped by processes operating in the vector. Alternatively, gametocyte timing might be selected for by within-host processes. For example, host immune responses that kill gametocytes of any age may be upregulated at night and so, parasites must compensate for this by maximising infectivity of the survivors at night. This assumes there are costs of being maximally infective throughout the circadian cycle and/or that gametocytes are constrained to only have a short period at maximal infectivity due to the processes involved in development and/or senescence. Testing for a role of host processes, such as rhythmic immune responses, requires separating the effects of parasite and host times-of-day on transmission. Perturbing parasite time independently of host time is challenging, but using circadian clock knockout mice would be a good start.

Our data reveal that mosquitoes fed at night are less susceptible to infection and that vector rhythms have long lasting effects on sporogony. Similarly, long lasting effects of time-of-day of infection are observed for the expulsion rate of the intestinal parasitic helminth, *Trichuris muris* [30]. Why might mosquitoes fed at night be less susceptible to infection? Rhythmic immune defences have been described in insects [31,32], including *Anopheline* mosquitoes [16]. For example, sporozoites are phagocytosed by haemocytes in mosquitoes [33] and the protective capacity of phagocytic cells peaks at night in *Drosophila* [31]. Furthermore, transcriptional profiling in *Anopheles gambiae* has revealed that at least 20 genes of known or putative immune function (including against malaria parasites) display diel expression patterns with variable timing (phases) [17]. Whatever the process(es) responsible, it appears they either protect mosquitoes from infection or not, rather than affect oocyst or sporozoite burden directly. That previous studies have not recognised the potential for time-of-day to affect the outcome of interactions between malaria parasite rhythms and vector rhythms, nor followed parasites throughout sporogony, may explain their lack of support for Hawking’s hypothesis.

Given that mosquito populations are responding to the use of bednets by shifting the time-of-day they forage for blood [18], understanding how time-of-day shapes interactions between parasites and mosquitoes is necessary to predict the epidemiological consequences of altered mosquito rhythms [34]. Clearly, if bed nets prevent night time transmission, then day-biting is beneficial for parasites, but if human malaria parasites behave as we observe for *P. chabaudi*, transmission potential may not change (*i.e.* mosquitoes fed at ZT8 on ZT8 parasites harbour the same sporozoite burdens as mosquitoes fed at ZT16 on ZT16 parasites). In contrast, if parasites can alter their rhythms, they may be able to capitalise on the greater transmission potential of (currently) night time gametocytes that infect mosquitoes feeding in the day. However, the fitness consequences of coordinating the development of asexually replicating stages with the host’s circadian rhythms [19,35] may constrain the capacity of parasites to adjust the timing of schizogony. In this case, the duration of gametocyte development may be selected on, necessitating investigations into the flexibility of gametocyte developmental duration.Furthermore, given that mosquitoes will be under selection to cope with blood feeding in the daytime by changing the timing of associated physiological processes [34], the overall impact on malaria transmission is hard to predict.

The notion that rhythms in transmission traits expressed by parasites maximise onwards transmission has also been applied to parasite species that do not require a vector. For example, peak shedding of *Isospora* sp. from its avian host occurs in the late afternoon and this timing is thought to minimise damage from UV radiation while the parasite waits to encounter a new host [36]. The cercariae of *Schistosoma* coincide are proposed to time their emergence from the intermediate snail host in the morning or afternoon according to whether they specialise on livestock or nocturnal rodents for their final host [37–39]. Finding support for Hawking’s hypothesis in malaria parasites should motivate tests of whether periodicity in transmission behaviours maximises fitness for other parasite lifestyles.

## Materials and Methods

### Ethics Statement

All animal procedures were carried out in accordance with the terms of the UK Animals (Scientific Procedures) Act (PPL 70/8546) and have been subject to ethical review.

### Hosts, parasites and vectors

Vertebrate hosts were 8 to 11 week old C57Bl/6J female mice given access to food and drinking water (supplemented with 0.05% para-amino benzoic acid [40]) *ad libitum*. We entrained forty mice to a 12:12 hrs light:dark cycle 3 weeks prior to and during the experiment. At ZT3, we infected all mice with 10^5^ rodent malaria *Plasmodium chabaudi* genotype AS (passage number A12) infected red blood cells at ring stage by intraperitoneal injection. ZT3 refers to 3 hours after lights on and we infected all experimental mice with parasites from donor mice entrained to the same photoperiod to avoid costs of mismatching the phase of parasite and host rhythms [19].

Mosquitoes were reared according to Spence *et al.* [41]. Briefly, we housed mosquitoes at 26°C, 70% relative humidity, 12:12 light:dark cycle, and provided them with 10% fructose solution, supplemented with 0.05% para-aminobenzoic acid. We randomly assigned *Anopheles stephensi* mosquito pupae to three offset photoschedules according to their treatment groups (SI Figure 2). At 5 days post pupation, we randomly assigned female mosquitoes within each photoschedule to paper cages with meshed lids (20 replicate cages for each of the four experimental groups, with 60 mosquitoes per cage) and supplemented their sugar/ para-aminobenzoic acid water with 0.05% gentamicin solution prior to blood feeding. We starved mosquitoes for exactly 24 hours before blood-feeding by providing them with access to water only. Blood feeds occurred when the mosquitoes were between 5-11 days post emergence, and mosquitoes were given sugar/ para-aminobenzoic acid water immediately following blood feeding.

### Experimental design and transmissions

On day 14 post infection, at ZT8 or ZT16, we took blood samples from all mice by tail snip for quantification of gametocytes by RNA extraction [42] and subsequent quantitative reverse transcriptase PCR (RT-qPCR) targeting the sexual stage-specific expressed gene PCHAS_0620900, previously named PC302249.00.0 [43]. Immediately after sampling, we anaesthetised mice as per PPL 70/8546 using an injection of ketamine hydrochloride and medetomidine, and exposed each mouse to its cage of mosquitoes for twenty minutes and then euthanised them, following Spence *et al.* [41]. We carried out all mosquito feeds in the dark to prevent differences in mosquito biting rates resulting from unexpected light exposure during blood feeding [21,22]. We performed the experiment in 2 identical blocks (n=10 feeds per treatment group per block), initiated 2 days apart.

Our experimental design involved four treatment groups in which mosquitoes fed at their ZT8 or ZT16 on mice at their ZT8 or ZT16. We reared all experimental mice in the same photoschedule, with lights on at 1:00 GMT and off at 13:00 GMT. To cross factor “time zones” for parasites and vectors we entrained mosquitoes to three offset photoschedules where lights on and lights off occurred at different times with respect to GMT (though always with 12:12 light:dark photoperiods) (SI Figure 2). Mosquito photoschedule 1 provided mosquitoes at their ZT16 to feed on ZT8 mice. Mosquito photoschedule 2 provided mosquitoes experiencing their ZT8 to feed on mice experiencing ZT8 as well as mosquitoes experiencing their ZT16 to feed on mice experiencing ZT16. Mosquito photoschedule 3 provided mosquitoes at their ZT8 to feed on mice experiencing ZT16. This design avoided the need to have multiple groups of mice, allowing all infections to be initiated from the same parasite stock and donor mice, whilst accounting for the phase-mismatched treatments being 8 hours apart.

We quantified infection in mosquitoes at oocyst (day 8 post blood meal) and sporozoite (day 14 post blood meal) stages. For oocyst quantification, we dissected midguts from 20 cold anaesthetized mosquitoes per cage and visualised them microscopically after staining with 0.5% mercurochrome. To ensure that all 800 mosquitoes could be processed within a single day, we took photographs of all midguts (*e.g.* SI Figure 3) and quantified oocysts using ImageJ software v1.51o (NIH, Bethesda, MD, USA). We quantified sporozoites from pools of 5 mosquitoes (n=4 pools per cage) that were bisected between the second and third pair of legs following Foley *et al*. [44], ensuring that salivary gland sporozoites are quantified, not those remaining in oocysts. We extracted DNA using the CTAB-protocol from Chen *et al.* [45] with minor modifications: tissue lysis was done by shaking a ball bearing at 30bpm for 2 minutes in a tissue lysate machine; liquid volumes were increased; a chloroform step and 10 minute chilled ethanol incubation were added; and all centrifuge steps were performed at 4°C. We used qPCR targeting the 18S rRNA gene [46] to quantify the number of parasite genomes, assuming 1 genome per sporozoite. Sporozoite data for one mouse (mouse ZT16/mosquito ZT8) were lost due to a failed DNA extraction.

### Additional experiments

We analysed an independently collected dataset to verify the observation of lower densities of gametocytes in host blood at night. Four days after administration of 125mg/kg phenylhydrazine to induce anaemia and enhance gametocyte production, four 8-week old C57Bl/6J male mice were intravenously infected at ZT2 with 10^7^ *Plasmodium chabaudi* genotype CR infected red blood cells at ring stage. On day 4 post infection, gametocyte densities were quantified at ZT5, 9, 17 and 21 by microscopy.

To investigate whether lower densities of gametocytes at night can be explained by gametocytes accumulating in the peripheral capillaries, we carried out an additional experiment. We used mosquitoes to sample blood from peripheral capillaries of mice and compared the density of gametocytes in their guts to those in the tail vein of the mice they fed on. Three mice were infected with 10^5^ *P. chabaudi* genotype ER infected red blood cells at ring stage. On day 14 post infection, we made thin blood smears for gametocyte quantification from the tail vein in triplicate, and immediately offered *An. stephensi* mosquitoes a blood meal following the same protocols as in the main experiment. We carried feeds out at ZT16 for both parasites and mosquitoes. Within 10 minutes post blood meal, we dissected 5 mosquitoes per feed and used the contents of their midguts to make thin blood smears. We stained all smears with Giemsa and quantified gametocytes by microscopy for the smears of tail vein blood (3 per mouse) and mosquito midguts (5 per mouse). Because one mouse did not have any gametocytes, its data were excluded from analysis.

### Statistical Analysis

We performed all data analyses using R v3.2.4 (R Foundation for Statistical Computing, Vienna, Austria). Gametocyte, oocyst and sporozoite densities were analysed using linear models, with a ^10^log transformation for gametocyte densities, and a square root transformation for oocyst and sporozoite burdens to meet assumptions of normality and homogeneity of variance. We used generalised linear models, with a binomial error structure, to analyse the proportion of fed mosquitoes and the proportion of infected mosquitoes (a two-vector binomial response variable including counts of fed / unfed or infected / uninfected mosquitoes). The proportion of sporozoite-infected PCR pools was analysed using generalised linear mixed models, with a binomial error structure, using mouse as a random effect to account for multiple PCR pools taken from the same mosquito feed. Gametocyte densities, normalised to WBCs, were compared between tail vein and mosquito midguts using linear mixed effect models with mouse fitted as a random effect. We minimised models after comparison with Log Likelihood Ratio Tests, for which test statistics and p-values are reported, to determine whether terms could be removed. The identity of each block was controlled for in all analyses. Block did not interact with any of the fixed effects fitted in any analyses and so, statistics for terms involving block are not presented in the results. However, sporozoite prevalence (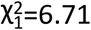, *P*=0.010) and sporozoite burden differed between the blocks (F_1,73_=4.89, *P*=0.030) and so, block is controlled for in the statistics presented for these analyses, with average effect sizes and SEM calculated across blocks.

## Acknowledgements

We thank Ronnie Mooney for technical assistance and Phil Spence, Wiebke Nahrendorf, Daan van der Veen, and Giles K.P. Bara for advice.

The work was supported by NERC and BBSRC (NE/K006029/1), the Royal Society (UF110155; NF140517), the Wellcome Trust (202769/Z/16/Z; PhD programme in Hosts, Pathogens and Global Health), and the Human Frontiers Science Program (RGP0046/2013).

## Author Contributions

PS, SSCR, KP, and SER conceived and designed the project. PS, NS, and AOD performed the experiments and data analysis. PS, SSCR, and SR wrote an initial draft of the manuscript. All authors contributed to interpretation of results and refinement of the manuscript.

